# Scrambling the skin: Simulated Skin Re-Arrangement Using Apparent Motion

**DOI:** 10.1101/2019.12.20.884239

**Authors:** Tatjana Seizova-Cajic, Sandra Ludvigsson, Birger Sourander, Melinda Popov, Janet L Taylor

## Abstract

An age-old hypothesis proposes that object motion across the receptor surface organizes sensory maps (Lotze, 19^th^ century). Skin patches learn their relative positions from the order in which they are stimulated during motion events. We test this idea by reversing the local motion within a 6-point apparent motion sequence along the forearm. In the ‘Scrambled’ sequence, two middle locations were touched in reversed order (1-2-<u>4-3</u>-5-6, followed by 6-5-<u>3-4</u>-2-1, in a continuous loop). This created a local acceleration, a double U-turn, within an otherwise constant-velocity motion, as if the physical location of skin patches 3 and 4 was surgically swapped. The control condition, ‘Orderly’, proceeded at constant velocity at inter-stimulus onset interval (ISOI) of 120 ms. In the test, our twenty participants reported motion direction between the two middle tactors, presented on their own at 75, 120 or 190-ms ISOI. Results show degraded motion discrimination following exposure to Scrambled pattern: for the 120-ms test stimulus, it was 0.31 d’ weaker than following Orderly conditioning (p = .007). This is the aftereffect we expected; its maximal expression would be a complete reversal in perceived motion direction between locations 3 and 4 for either motion direction. We propose that the somatosensory system was beginning to ‘correct’ reversed local motion to uncurl and remove the U-turns that always occurred on the same part of the receptor surface. Such de-correlation between accelerations and their location on the sensory surface is one possible mechanism for organization of sensory maps.

## II. INTRODUCTION

*“When, in movement of the body, a stimulus changes its region of stimulation, the local signs change, and successive local signs are the things of adjacent localities” (19th century philosopher Lotze, cited in [1], p. 268)*

Somatosensory projection areas in the brain are dubbed brain maps because they reflect the topographical layout of the receptor surface. How maps develop and remain calibrated throughout life is a question of long-standing interest. It was empirically addressed in the classical study on synaptic plasticity by Merzenich and Jenkins [2]. In monkeys, they performed an anatomical (surgical) manipulation of the receptor surface by relocating a flap of skin from digit 4 to digit 3, fully preserving all its original innervations. Several months after the transfer, stimulation of the relocated flap excited cells in the cortical area that previously only represented the finger to which the flap was relocated. Since the surgery created new patterns of co-stimulation of different skin parts, the authors concluded that cortical representations are time-coincidence-based concepts.

In subsequent research, timing has usually been conceptualized and operationalized as temporal coincidence or neural co-activation (see [3, 4, 5]), although motion across the receptor surface is a better candidate for the general organizing principle of spatial maps. It is a ubiquitous form of natural stimulation and, importantly, unlike simultaneous (coincident) stimulation, it cannot lead to ‘fusion’ of skin parts that often touch each other, such as lips or fingers. We therefore revive an old hypothesis (one source is quoted above) that *motion* organizes spatial maps in touch and vision. (There is of course a significant genetic component to topography, evident during embryonic development, but it is crude and insufficient [6]. There is also spontaneous, synchronized oscillatory electrical activity independent of interactions with the world, which occurs prenatally and in early development, when plasticity is high (in rodents and humans [6, 7]).)

Motion is a very strong candidate for the experience-dependent map organization because locations next to each other on a sensory surface are stimulated one after the other by moving objects, and can thus learn that they are neighbours. The idea is illustrated in Fig 1 (and was previously described in [8, 9]). Although quite simple and old (Lotze, 19^th^ C, cited in Herrnstein and Boring, 1965), it has attracted surprisingly little research and has little direct evidence to support it (to our knowledge, none in humans). Motion was used in two animal studies and it strongly influenced brain maps. One was a vision study, in which investigators reversed direction of optic flow in young tadpoles before the map was developed, resulting in a poor retinotopy. They concluded that *“visual information is transformed from a temporal code to a spatial code in the brain”* (p. 1, [10]). In another study, [11], a rectangular flap of skin on a belly of the rat was rotated by 180 degrees, preserving the innervation as in the Merzenich study described earlier. One group of animals was subsequently exposed to brushing stimulation across the line of skin incision for 7 hours, while control animals received no such input. The cortical neurons in the experimental group developed significant changes in receptive fields consistent with the new skin arrangement, unlike the control group, demonstrating that motion stimulation merely hours long is an effective stimulus for such a change.

**Fig. 1.**
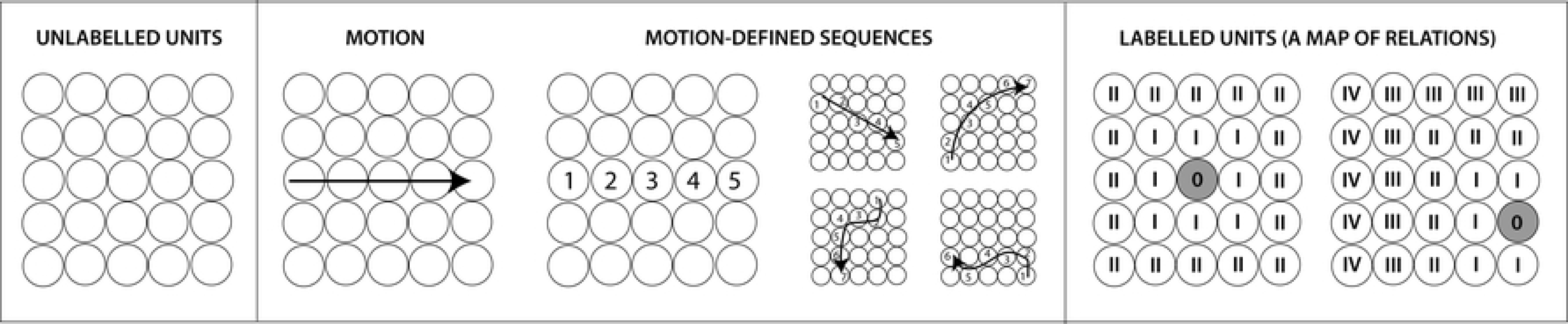
An illustration of the idea that motion across a sensory surface informs about neighbourhood relationships. Left panel: Units in a 2-D array (representing sensory neurons) have no ‘labels’ indicating their position in the array. Middle panel: Object motion proceeds in a sequence, activating units along its trajectory. Exposed to numerous motion events, adjacent localities will often be stimulated one after the other, as indicated by numerical sequences here. Right panel: As the outcome of this stimulation, units gain ‘labels’, i.e., the system gains information about their relative position in the array. Consider the central unit (labelled ‘0’ in the picture on the left): its first-degree neighbours are the units stimulated immediately before or after (labelled ‘I’); its second-degree neighbours (‘II’) are adjacent to its first-degree neighbours, etc. Each unit has a neighbourhood network (see a different unit depicted in the image on the right). Combination of these relationships makes a spatial map i.e., an array able to distinguish between different spatial configurations impinging on it.

The latter study is conceptually related to the our present study, because we attempted to *simulate* skin re-arrangement. We have done so in conscious humans, using apparent (sampled) motion. Our aim is to provide support for the idea presented above, that any two locations are assigned their relative positions based on the order in which they are stimulated during object motion. By definition, the order of skin stimulation is consistent with motion trajectory: an object moving in a proximal direction – for example, up the forearm - will stimulate a more distal location before its proximal neighbor (Fig 2, left). However, by using discrete stimuli to produce apparent motion it is possible to reverse this order for two skin patches in the middle of the motion trajectory. That is, during proximal motion, a more proximal patch can be stimulated before its distal neighbor (Fig 2, right, locations 4 and 3, respectively), and vice versa. We repeatedly applied such a scrambled motion pattern to the forearms of human participants to determine whether it could alter their perception of relative locations on the skin. Repeated many times, in exactly the same location, we expected this pattern to result in subsequent perceptual error consistent with the swapping of places for affected locations (locations 3 and 4 in Fig 2, right). Such an error could be revealed using a task that relies on position coding of the two locations. The experiment below tests this prediction using a two-point motion as a test stimulus.

**Fig 2.**
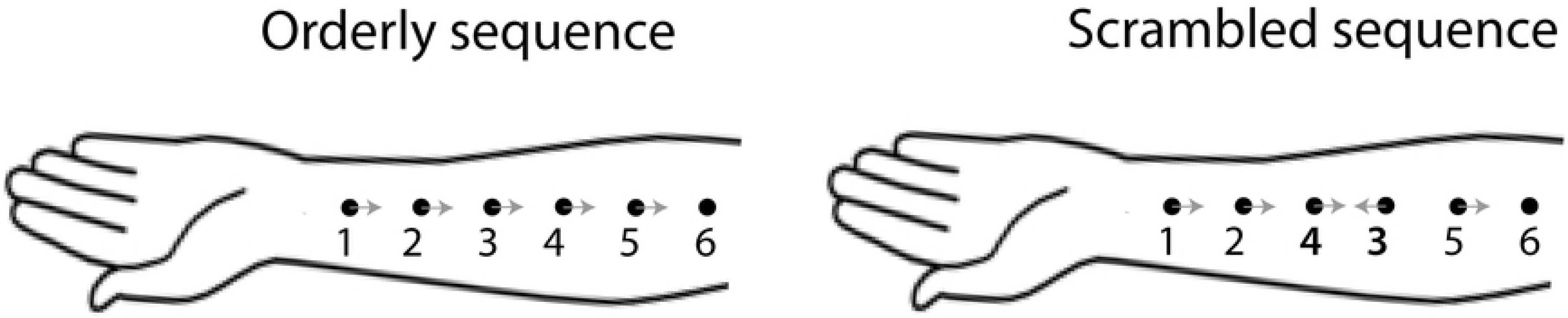
Orderly and scrambled patterns of apparent motion across the skin. Dots indicate touched locations. In both patterns, motion begins near the wrist and finishes near the elbow. They differ in the order of stimulation of the middle two skin patches only, as indicated by numbers. Grey arrows indicate direction of local stimulus motion i.e., motion between sequential stimulus pairs. Local motion has opposite direction to global motion in the middle of the motion trajectory in the Scrambled sequence.

## III. METHOD

### A. Participants

Twenty volunteers participated in the study (age range 18-30, 12 females), which was approved by the University of Sydney Ethics Committee. They were all naïve regarding study aims and design, and were paid $20 per hour. All participants provided written consent prior to participation.

### B. Overview and study design

The experiment tested the ability to judge motion direction of a *test* stimulus following exposure to a *conditioning* stimulus. Study design of this repeated-measures experiment is described in Figs. 3 and 4.

**Fig. 3.**
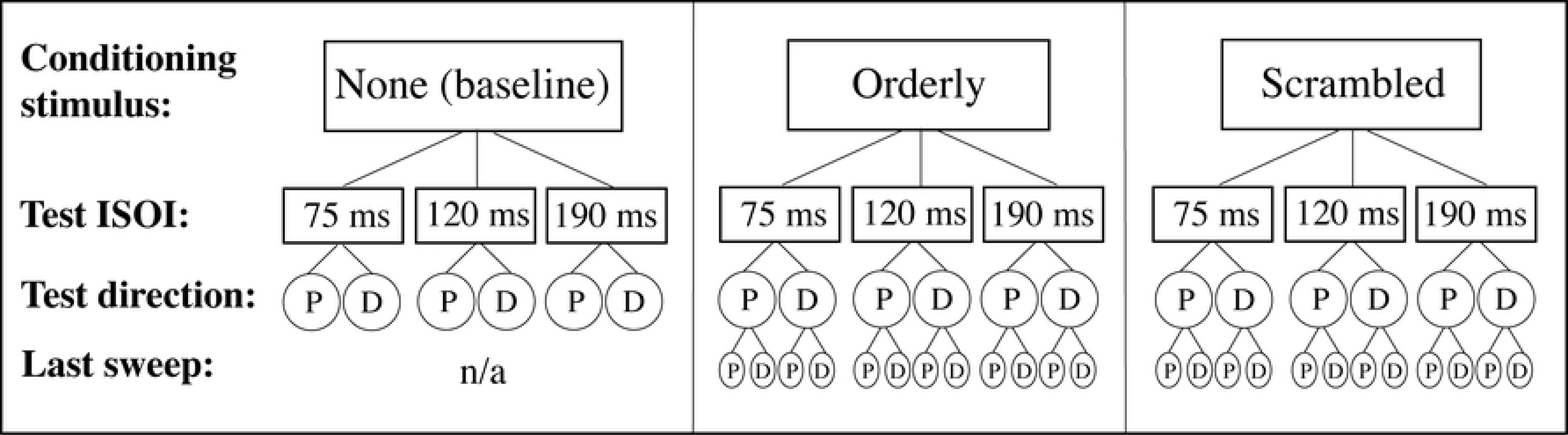
Experiment design. ISOI stands for Inter-Stimulus Onset Interval, P for proximal motion, and D for distal motion. ‘Last sweep’ refers to the last sweep in the conditioning sequence. See text for more details.

**Fig 4.**
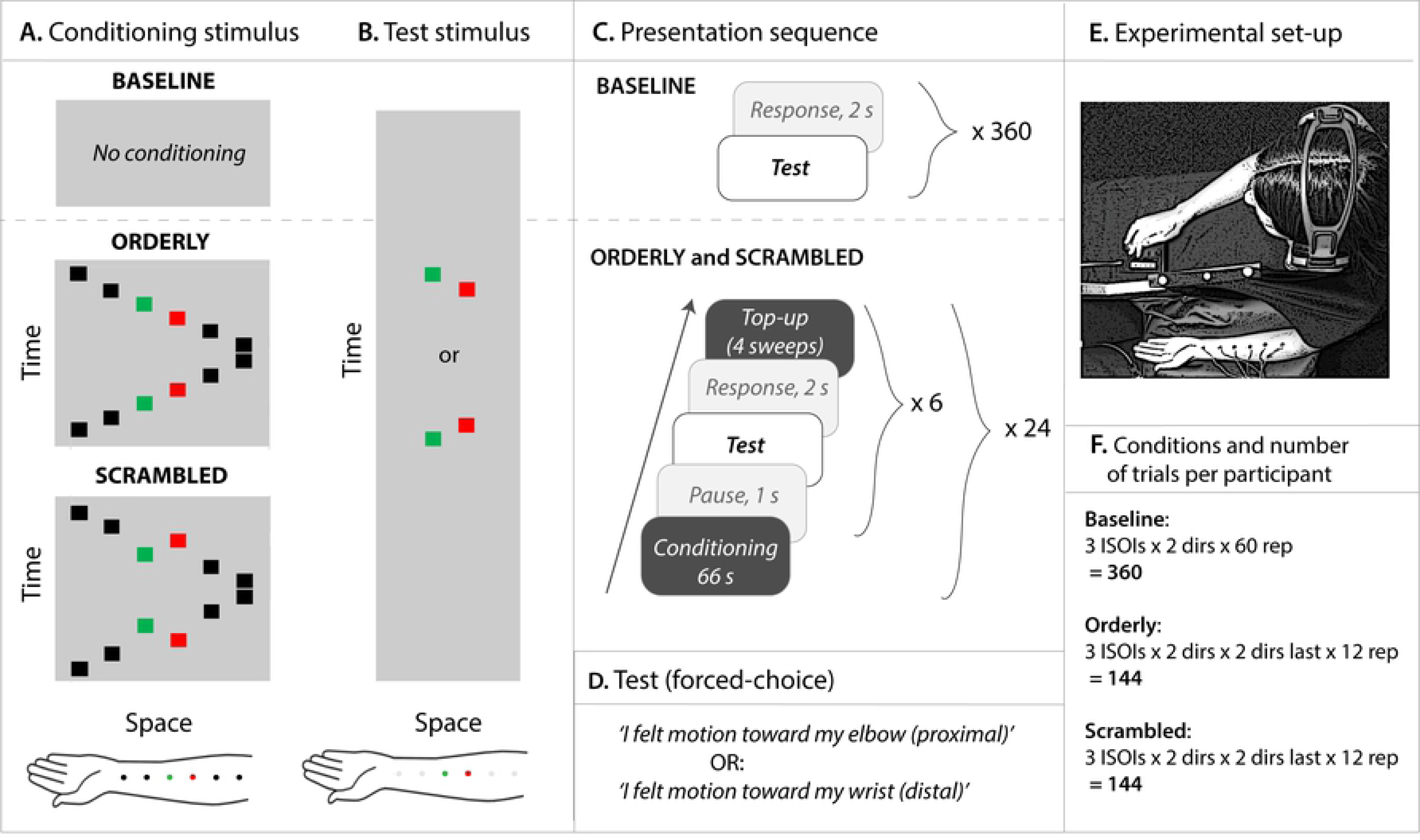
Details of the method. **A.** Conditioning was used in the Orderly and Scrambled conditions, shown here as space-time diagrams. Time in arbitrary units is represented on the Y-axis, and space (along the forearm) on the X-axis. Duration of one back-and-forth sweep was approximately 1440 ms. The black and coloured squares represent position of vibrators used in these conditions. **B.** Test stimulus, presented as time-space diagram, was the same in all conditions. The coloured squares represent vibrators used in these conditions and grey squares, vibrators attached to the forearm but not used. **C.** Stimulus sequence consisted of conditioning, 1-s break, test, 2-s for response, top-up, followed by five more repeats of the test-response-top-up cycle. **D.** Forced-choice task used to judge motion direction in the test stimulus. **E.** Bird’s eye view of the experimental setup. Six vibrators were attached to participant’s left forearm throughout the experiment, occluded from participant’s view. White noise presented through headphones masked the sound of vibrators. The participant responded to test stimuli by pressing one of the two buttons on the response box. **F.** Conditions and number of trials per participant (‘dir’ stands for direction of the test stimulus, ‘dir last’ for the direction of the last conditioning sweep, and ‘rep’ for the number of repeats per condition). Vibrator placement, our second control variable, is not explicitly presented for simplicity; it is embedded in repeats, such that each of the vibrator orders was used in half of the repeats.

The participant’s task was to report direction of apparent motion for stimuli applied by a pair of tactors. The direction was either proximal or distal. Perceived direction of the test stimulus obtained from this forced-choice task was the dependent variable. The main independent variable was the conditioning stimulation pattern (top row of Fig. 3). In **Baseline**, these test stimuli were presented with no prior conditioning (Fig. 4A, *Top left panel*). In the control condition, **Orderly** (Fig. 2A; Fig. 4A, *Middle panel*), the conditioning motion sequence proceeded across the skin at constant velocity (1-2-3- 4-5-6, or 6-5-4-3-2-1). In the **Scrambled** condition (Fig. 2B; Fig. 4A, *Bottom panel*), the activation sequence for locations 3 and 4 was reversed (1-2-**4-3**-5-6-5-**3-4**-2-1). The inter-stimulus onset interval (ISOI) in the conditioning sequence was always 120 ms, and the inter-stimulus interval (ISI) was 0 ms.

Following conditioning with either pattern, test stimuli comprising locations 3 and 4 were presented in the order 3-4 or 4-3. The second independent variable was Inter-Stimulus Onset Interval (ISOI) for the test stimulus (second row of Fig. 3). Three values chosen after extensive piloting were **75, 120** and **190 ms**, while piloting itself was informed by the literature (see [12, 13]). Our intention was to include stimuli difficult to discriminate - as was the case for the ISOI of 75 ms, and those relatively easily to discriminate (190 ms ISOI). Duration of vibration at each skin location was equal to the ISOI. For example, in the 75 ms condition, each vibration lasted 75 ms, immediately followed by vibration at the next location. Pilot studies suggested that zero-ISI results in the smoothest perception of motion. The third independent variable was direction of motion in the test stimulus: **proximal** or **distal** (third row of Fig. 3; Fig. 4B).

We controlled for the direction of the last sweep in the conditioning stimulation (fourth row of Fig. 3). Motion direction in the conditioning stimulation alternated: each sweep in one direction was followed by the sweep in the opposite direction. This should have created equal net adaptation and no net directional aftereffect (directional aftereffect in tactile motion is the bias to perceive direction opposite to the preceding motion; see [14, 15]), except for the possible greater influence of the last sweep in the conditioning stimulus, which was followed by the test. To control for the direction of the last sweep, half of conditioning trials ended with a proximal sweep, randomly interleaved with distal last sweeps.

The final variable we manipulated (the second control variable) was placement of the vibrator array (not shown in Fig. 3). Vibrators were numbered 1-6, and in one half of each participant’s session, they were physically placed in order 1-6 proximo-distally (from near the elbow crease toward the wrist), and in the other half, in reversed order (6-1). Which order was used in which half-session was counterbalanced across subjects.

The total number of different experimental conditions was 30: 6 in Baseline (3 ISOIs x 2 directions of test motion) and 12 each in Orderly and Scrambled (3 ISOIs x 2 directions of test motion x 2 directions of the last conditioning sweep). Perception of motion direction in the test stimulus was assessed using a forced-choice task.

### C. Apparatus, set up and procedure

Baseline, Orderly and Scrambled stimuli were presented in separate sessions, on separate days, in a partially counterbalanced order across participants (counterbalancing was imperfect because we had 20 participants and could thus not have an equal number of all possible orders of the three conditions). Presentation sequences are illustrated in Fig. 4C. No conditioning stimulus was presented in the Baseline condition, and test stimuli were presented one after another, separated by a two second break for response, divided into two equal blocks of 180 stimuli with a break in-between. In the Orderly and Scrambled conditions, participants initially received 66 seconds of conditioning with stimulus motion up and down the forearm. After a one-second break, they were presented with one test stimulus and had two seconds to report direction of apparent motion in the test stimulus (‘proximal’ or ‘distal’, see Fig. 4D) using a response box. Immediately afterward, they received top-up conditioning consisting of two sweeps up and down the forearm, followed by another test. Six test stimuli were presented in this manner, separated by top-ups. After a short break, the whole cycle was repeated (66 s of conditioning, and six test-response-top-up sequences). This was repeated 24 times in the Orderly and Scrambled sessions, divided into two blocks of 12 each, with a 10-minute break in between. Placement of the vibrator array (the first control variable described earlier) was different in the two blocks.

The radial aspect of the forearm, hidden from participant’s view (see Fig. 4E), had a linear array of 6 coin-motor vibrators attached to it, placed 4 cm apart, centre-to-centre. Activating vibrators one after another created perception of apparent motion. The ISOI in the conditioning stimuli was 120 ms, equal to duration of vibration. One sweep up or down the forearm lasted 720 ms. The test stimulus was presented in the order 3-4 or 4-3, at one of the following ISOI: 75 ms, 120 ms and 190 ms, (corresponding to the velocities of 53.3, 33.3 and 21.1 cm per second, respectively).

The vibrators used to create a sense of apparent motion were 10 mm in diameter, 3 mm high cylindrical coin motors (Precision Microdrive^TM^), in which eccentric rotating mass results in vibration. They were controlled by by a custom developed software (LabView^TM^ 2012). We used a laser apparatus (OptocoNCDT 2200; data extracted using LabChart^TM^) to measure vibration frequency and the degree of vibration transmission to the surrounding skin. Vibration frequency was initially unequal for different vibrators, but adjustment of current brought them all to approximately 110 Hz (to control for any remaining differences, the order of vibrators was reversed in half of the trials, as described earlier). Laser measurements also showed that vibration transmission via the skin occurs over at least 4 cm distance from the vibrator. This was consistent with a perception test, in which a fingertip was placed at different distances from the vibrator attached to another person’s forearm. Skin vibration was in some instances detectable 8 cm away from the vibrator, double the 4-cm separation between the vibrators we used. Thus, the stimulus delivered to a particular location affected a much greater area, adding noise to our desired spatiotemporal stimulus pattern (but we don’t know how much of the vibration spread was above threshold).

As Fig. 4F shows, the total number of trials in Baseline was 360 (3 ISOIs x 2 directions x 60 repeats), and in Orderly or Scrambled, 144 (3 ISOIs x 2 directions x 2 directions of the last sweep x 12 repeats). Each participant thus completed a total of 648 trials.

Each session was preceded by a short practice, which differed between conditions. In the practice for the Baseline condition, 90 test stimuli were divided into three blocks separated by two short breaks. Practices for Scrambled and Orderly conditions began with 60 test stimuli, followed by three bouts of conditioning containing 6 tests each.

A short questionnaire was used at the end of Scrambling and Orderly sessions to capture participants’ perception of the 66-s conditioning stimulus. The participants ranked the frequency of experience on a 7-point scale ranging from ‘never’ to ‘always’. Example statements are: *During the longer (1-min) periods of stimulation, I felt…‘Motion on my forearm’; ‘Motion along the straight line’; ‘Curved or zig-zag motion path’*. The order of questions was randomized for each participant.

Participants also answered two open-ended questions and gave a phenomenological report by sketching what they had felt on a standardized picture of an arm.

### D. Data analysis

Raw data were responses regarding the direction of motion (distal vs proximal) of the test stimuli. R (R Foundation, R 3.1.0, 2014) was used to extract data and compute the proportion of correct responses for each participant in each condition, and SPSS for further analysis. There were 12,960 possible responses (20 participants x 648 responses), of which 217 or 1.7% were missing (failed to record – either the participant did not respond, or they pressed the response button with insufficient force).

We used signal detection theory to compute sensitivity to motion direction (d’) and bias (c). This absorbed one of our independent variables, direction of test motion. D-prime was defined as the difference between z-scores for the proportion of correct responses to proximal test motion and the proportion incorrect to distal test motion. Bias was the average of the same pair of z scores, multiplied by −1 (‘c’ measures of bias, [16], p .143). Computed in this manner, negative c means bias toward the response ‘proximal’.

The example in Table 1 shows steps in computing d’ and c for one experimental condition (Scrambled, 120 ms SOA) in one participant. Note that the four values each of d’ and c (shown in the last two rows of Table 1) were computed based on a total of 48 stimuli presented, 6 repeats for each of the 8 conditions specified in the table (2 directions of test motion x 2 Vibrator placement options x 2 Directions of the last conditioning sweep). All four values were used in linear mixed modelling (we did not average them) to allow us to test for quadratic trends across the three SOAs. Variability between the 4 measures of d’ and c included variations, if any, caused by our two control variables (Vibrator placement and Direction of the last sweep).

**Table 1.** Example computation of d’ and c, Scrambled condition, 120-ms SOA, one participant.

|  | Test stimulus motion (6 stimuli per condition) |  |  |  |  |  |  |  |
| --- | --- | --- | --- | --- | --- | --- | --- | --- |
|  | Proximal |  |  |  | Distal |  |  |  |
| Vibrators 1-6 placement | Elbow to wrist |  | Wrist to elbow |  | Elbow to wrist |  | Wrist to elbow |  |
| Last conditioning sweep | dist. | prox. | dist. | prox. | dist. | prox. | dist. | prox. |
| Correct response | 6 | 4 | 5 | 3* | Redundant (total responses per condition = 6) |  |  |  |
| Incorrect response | Redundant (total responses per condition = 6) |  |  |  | 4 | 3 | 1 | 0 |
| Hit rate for <u>Proximal</u> motion | 1.00 | 0.67 | 0.83 | 0.60 |  |  |  |  |
| False Alarm (FA) rate for <u>Proximal</u> motion |  |  |  |  | 0.67 | 0.50 | 0.17 | 0.00 |
| Hit rate for <u>Proximal</u> motion, corrected (0 becomes 0.05; 1 becomes 0.95) | 0.95 | 0.67 | 0.83 | 0.60 |  |  |  |  |
| FA rate for <u>Proximal</u> motion, corrected (0 becomes 0.05; 1 becomes 0.95) |  |  |  |  | 0.67 | 0.50 | 0.17 | 0.05 |
| Z score for corrected Hit rate | 1.64 | 0.43 | 0.97 | 0.25 |  |  |  |  |
| Z score for corrected FA rate |  |  |  |  | 0.43 | 0.00 | -0.97 | -1.64 |
| Ability to discriminate proximal from distal test motion ($d' = z_H - z_{FA}$) | 1.21 | 0.43 | 1.93 | 1.90 | | | | |
| Bias toward response 'distal' ($c = -(z_H + z_{FA})/2$) | -1.04 | -0.22 | 0.00 | 0.70 | | | | |

\*One response was missing in this condition; thus $3/5 = 0.60$ hit rate

A further 30 responses were excluded (single participant data, Baseline, 120 SOA, Vibrator placement 1-6) because d’ and c computed from them were both outliers. The corresponding Scrambled and Orderly conditions were also excluded.

To account for change in direction discrimination due to adaptation from the conditioning, we compared Baseline with the other two conditions. Our main question was addressed by the sensitivity analysis: we expected conditioning with Scrambled motion to result in reduced sensitivity (more motion reversals) compared to Orderly. For this analysis, we subtracted d’ in Baseline from each of the other two conditions, and compared Orderly and Scrambled to each other only. We analyzed response bias in a similar manner.

Linear Mixed Modelling (LMM) for repeated measures data [17] was performed via GLM procedure in SPSS. LMM accounts for the repeated nature of the data and for random variation across individuals. It also allowed the independent variable ISOI in the test stimulus to be treated as a continuous measure. Fixed factors were Conditioning pattern and ISOI. Participants were treated as a random factor, removing a significant proportion of within-subject covariance from the residuals; the Repeated subcommand in LMM dealt with the remaining deviations from the assumptions of a linear model. Both d’ and c were analyzed in this manner.

Questionnaire results were expressed on 0 (‘Never’) to 6 (‘Always’) scale, and summary measures were compared across conditions.

## IV. RESULTS

### A. Sensitivity to motion direction

Detailed results for Baseline, Orderly and Scrambling conditions are shown as box plots in Fig 5. Data files are available at the Open Science Framework <u>data repository</u>. The ability to discriminate direction was highest in Baseline (white boxes in Fig 5, *Left panel*). It increased with ISOI, approaching the ceiling at 190 ms. Sensitivity was lower both in Orderly (light grey) and Scrambled (dark grey) compared to Baseline.

**Fig 5.**
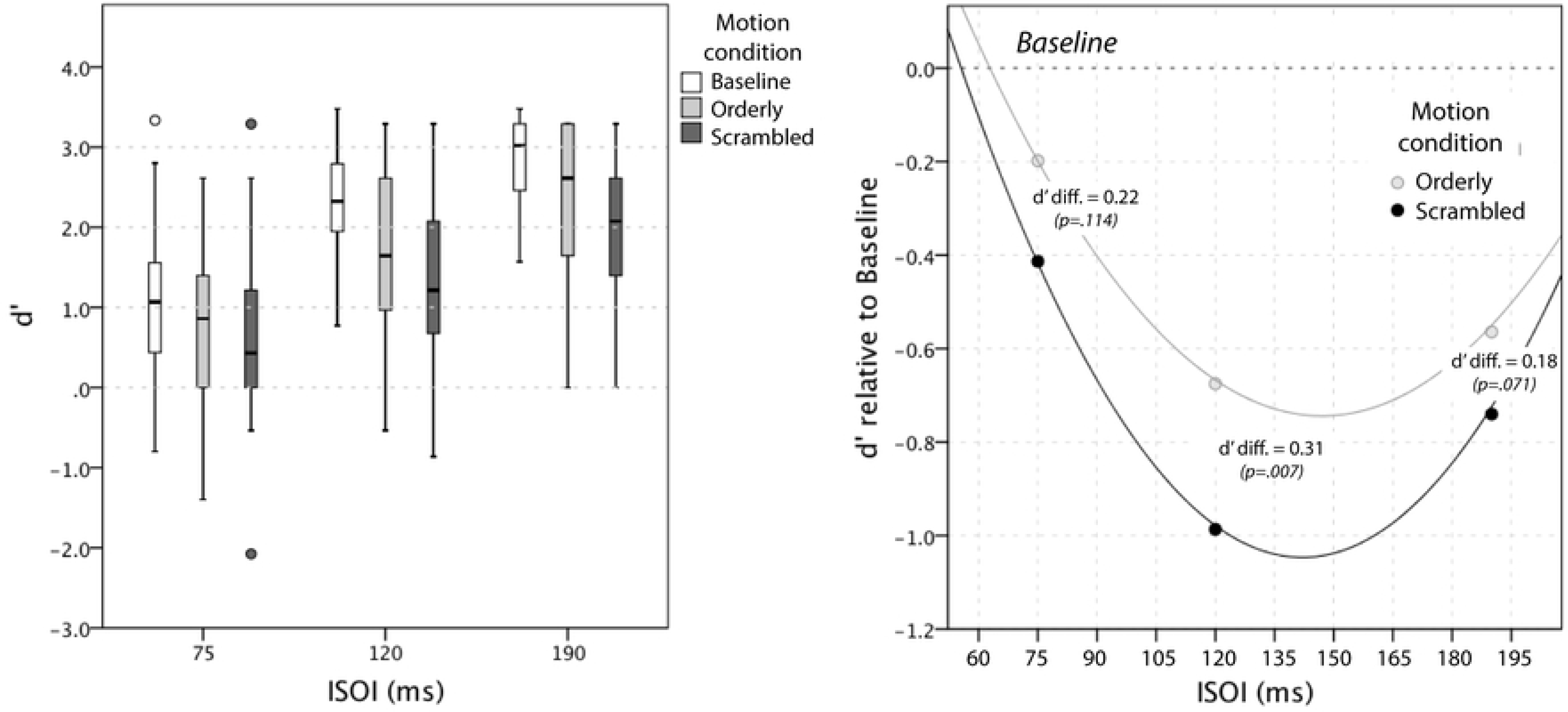
Sensitivity to motion direction, results. *Left panel*. Box plots show medians and variability in d’ for 20 participants as a function of ISOI and Motion condition. Note (a) the advantage of Baseline at all ISOIs, (b) the advantage of Orderly over Scrambled at al ISOIs, and (c) a ceiling effect at 190 ms, most pronounced for Baseline. *Right panel*. Estimated marginal means from LMM analysis of Baseline-corrected results (d-prime values are negative because Baseline was superior to both Orderly and Scrambled). The fact that Scrambled stimulation produced higher negative d’ values indicates worse discrimination of direction compared to Orderly. See text for details.

Our critical result is shown in Fig. 5, *Right panel*: reversed direction of the test stimulus (opposite to that actually presented) was perceived more frequently in Scrambled condition compared to Orderly. This is indicated by lower sensitivity (lower d’) in that condition. Note that d’ values in Fig 5, Right are negative because they are shown relative to Baseline – they simply show that performance was worse than in the Baseline (they do not indicate that perceived direction was reversed overall).

Linear mixed modelling (LMM) was used to estimate the difference in sensitivity (d’) between Orderly and Scrambled conditions after each of them was corrected for Baseline. Estimated quadratic functions are shown in Fig. 5, *Right panel*, and regression coefficients are given in Table 2. The effect of Conditioning pattern was significant (F(78.2, 1) = 7.82, p = .007), as was the ISOI (F(49.5, 1) = 10.68, p = .002). Sensitivity for motion direction was lower in Scrambled than Orderly condition. Compared to Baseline, it changed with ISOI following a quadratic trend (F(77.3, 1) = 15.15, p < .001), and the change was greatest for the ISOI of 120 ms. The interaction between Motion condition and ISOI was not significant (F(76.5, 1) = 0.06, p = .801), nor was it its interaction with ISOI^2^ (F(75.9, 1) = 0.86, p = .357).

**Table 2.** Results of linear mixed modelling (LMM), analyses of d’ and bias (c); see text for details.

| Analysis of $d'$ | | Analysis of bias | |
| --- | --- | --- | --- |
| Dependent variable | $d'$ | Dependent variable | c |
| *Fixed factors* | *Coefficients* | *Fixed factors* | *Coefficients* |
| Intercept | -0.97864 | Intercept | 0.1076 |
| ISOI (centered; 120 = 0) | -0.0062 | ISOI (centered; 120 = 0) | 0.00082 |
| Scrambled | reference | Scrambled | reference |
| Orderly | 0.31143 | Orderly | -0.01272 |
| Orderly x ISOI | 0.00049 | Orderly x ISOI | Tested and excluded from the model |
| Scrambled x ISOI | reference | Scrambled x ISOI |  |
| ISOI<sup>2</sup> | 0.00014 | ISOI<sup>2</sup> |  |
| Orderly x ISOI<sup>2</sup> | 0.000035 | Orderly x ISOI<sup>2</sup> |  |
| Scrambled x ISOI<sup>2</sup> | reference | Scrambled x ISOI<sup>2</sup> |  |

Estimated differences between d’ in Orderly and Scrambled from this model were 0.22, 0.31 and 0.18 at 75 ms, 120 ms, and 190 ms, respectively, with the following p values for Bonferroni-corrected pairwise comparisons: .114, .007 and .071.

### B. Bias in judgments of motion direction

Detailed results for the three motion conditions are shown in Fig 6. Summary of raw values of c is shown on the left, and absolute values on the right. Absolute values were computed because proximal and distal bias cancel each other out, potentially misrepresenting the strength of each individual’s bias and its variation across conditions.

**Fig 6.**
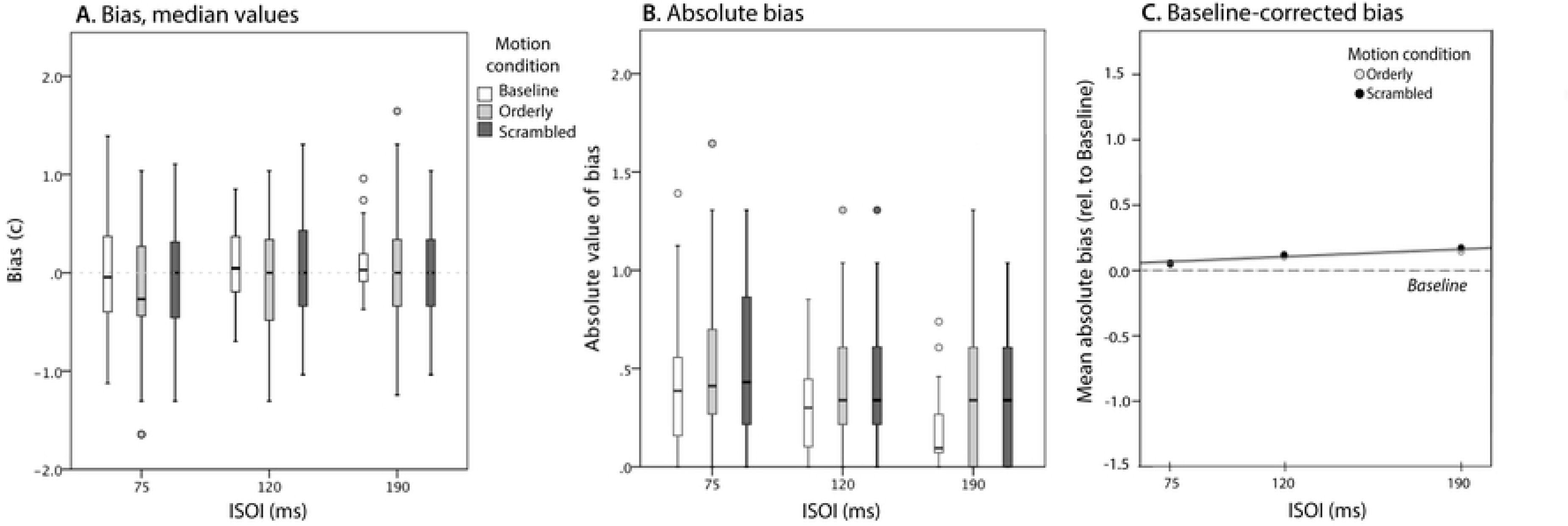
Bias in judgments of motion direction. **A.** Bias as a function of ISOI and Motion condition. Positive bias is tendency to report distal motion. Note that most medians are close to zero. **B.** Absolute values of bias, computed separately for each participant and condition. **C.** Baseline-corrected absolute bias for Orderly and Scrambled conditions, group means and linear functions estimated using linear mixed modelling. Note that the lines almost completely overlap, and that both Orderly and Scrambled conditions produced slightly more biased responses than Baseline.

There was almost no systematic bias at the group level (Fig. 6A), with little variation across conditions. Absolute values (Fig. 6B) show more bias and more variation. The condition with least absolute bias was Baseline at 190 ms (the stimulus easiest to judge – see Fig. 5, *Left panel*).

Baseline-corrected Orderly and Scrambled absolute bias is shown in Fig. 6C. Slightly greater than in Baseline (represented by a dotted line), bias is very similar in the two conditions: their means and linear functions estimated using LMM practically overlap. The effect of Motion condition was not statistically significant (F(1, 79.5)=0.226, p=.635). A mild increase in bias with ISOI estimated by the model (0.082 per 100 ms) was also not statistically significant (F(1, 18.2)=2.726, p = .116).

### C. Phenomenological reports

Answers to the questionnaire designed to explore perception of the conditioning stimuli are summarized in Fig. 7. It shows medians and standard errors for 19 participants (one participant’s data are missing due to experimenter error). Orderly and Scrambled conditioning stimuli were experienced similarly: all participants in both conditions felt motion up and down the forearm, mostly along the straight line, with occasional irregularities in the motion path (gaps, curves, zig-zag motion, twists and turns). Most of the time, it appeared to them that one object was moving, and sometimes two or more.

**Figure 7.**
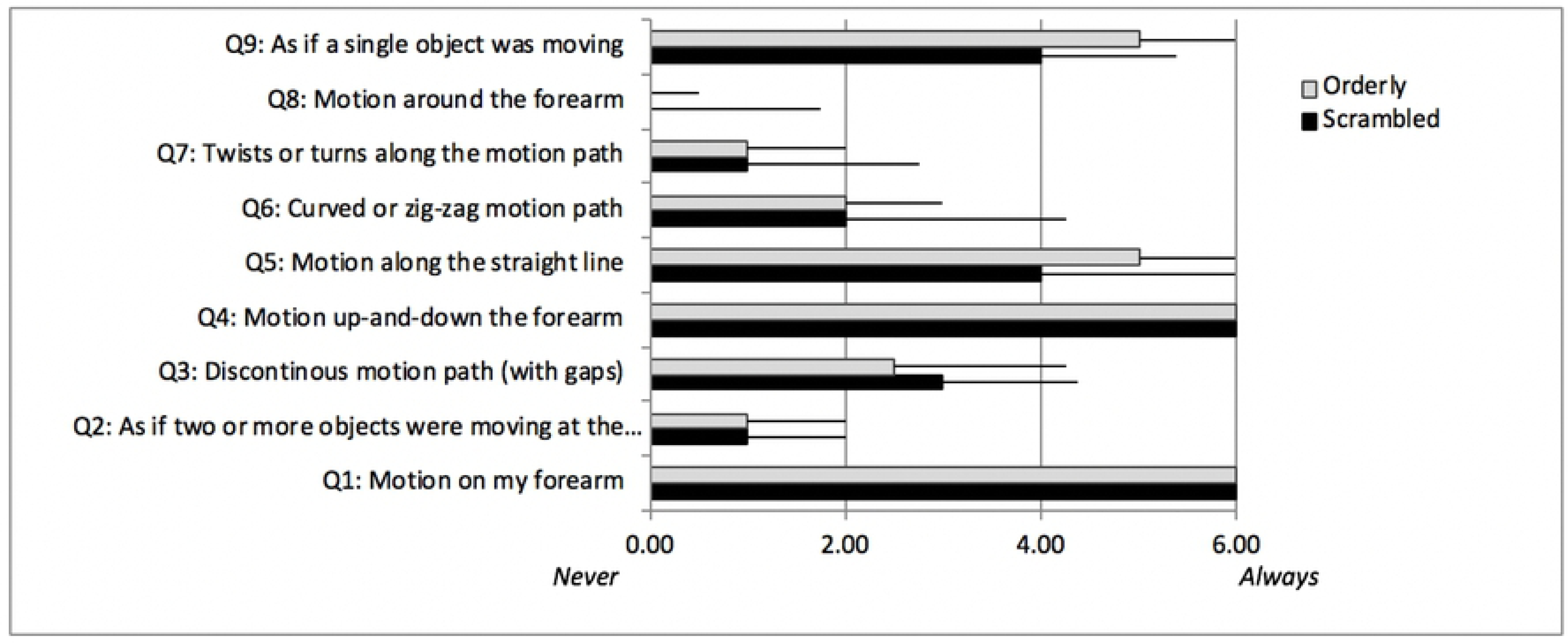
Answers to the questionnaire designed to explore perception of the conditioning stimuli (medians and standard errors, n=19). To ensure participants were referring to the conditioning stimulus rather than test stimulus, the root question asked: ‘*During the longer (1-min) period of stimulation, I felt…*’)

A one-point median difference was found for questions 5 (‘I felt motion along the straight line’) and 9 (‘I felt as if a single object was moving’), both more frequently experienced in the Orderly condition. Half a point median difference was found for question 3: ‘I felt a discontinuous motion path (with gaps)’, more frequently experienced in the Scrambled condition.

Drawings were scrutinized for any systematic differences between the two conditions, including presence of gaps and other irregularities, but there was no clear trend. All 20 pairs can be seen in the Open Science Framework <u>data repository</u> [https://osf.io/gtcr7/?view_only=87c6fb7b513b49758aa7185dcdf0e984].

## V. DISCUSSION

Sensory systems respond to their ‘diet’ (see [18] and [19] for an early and recent reference). Sufficient exposure to a new diet should result in a change, provided the system responds to the altered aspects of the diet. The sensory diet we provided was bi-directional apparent motion lasting 26.4 minutes in total (per session), created using discrete vibration stimuli and delivered in 66-s bouts of conditioning, interspersed with tests. In response, both scrambled and orderly conditioning was followed by a reduced ability to discriminate motion direction in the test stimulus relative to baseline. It was reduced at all test speeds, and mostly so for the 33 cm/s test, which matched the adapting speed (ISOI = 120 ms, see Fig. 5 *Right*). The crucial aspect of the diet we varied was the order of stimulation of skin patches. The scrambled sequence resulted in worse test performance than the orderly sequence, as predicted. The difference was again greatest for the 33 cm/s test.

Qualitative data (see Fig. 7) and the drawings show that the two conditioning patterns were similarly perceived: the participants felt motion up and down the forearm, mostly along the straight line. A variety of tactile and visual spatiotemporal patterns containing sudden accelerations are misperceived such that percept tends to be smoother than the stimulus patterns [8, 9, 20, 21]. Vision research also shows that transient changes in the motion sequence such as gaps in the trajectory or changes in colour or shape of the moving object are imperceptible provided they do not occur too early in the motion sequence [22].

Why were different results obtained in the orderly and scrambled conditions?

### A. Causes of different adapted states in the orderly and scrambled conditions

Intensity adaptation [23] and adaptation to motion [14, 15, 24, 25] can potentially both account for the impaired ability to distinguish proximal from distal motion in the orderly and scrambled conditions. Our main prediction was that conditioning with the scrambled pattern would cause an *even worse* performance in a subsequent test than the orderly pattern. This prediction was confirmed. We stimulated exactly the same skin locations an equal number of times in both conditions, therefore the intensity adaptation alone cannot explain the difference. The explanation must lie in the temporal sequence of stimulation. In the introduction, we argued that sequences of stimulation caused by object motion across the skin define relative positions of elements within a somatosensory map (illustrated in Fig. 1). In what follows we delve further into that explanation, followed by other potential explanations of the present results (not mutually exclusive).

### B. Adaptation as map change due to a new diet of motion patterns

The idea we explore is that elements in a map get assigned their relative positions based on the order in which a moving object stimulates them. We reversed motion direction over locations 3 and 4, creating a local motion opposite in direction to the global motion, as if the order of skin patches underneath vibrators 3 and 4 were actually swapped. We propose that an adaptive process began to *re-assign relative positions* of somatosensory neurons with receptive fields in locations 3 and 4 accordingly. A perceptual consequence of such a process should be reversal in perceived direction of motion across the two locations, consistent with our findings.

Is the proposed process feasible? Given that objects in the world often accelerate, if accelerations were to cause map change, a consequence of acceleration-triggered map changes could be instability in neural networks. However, this should not happen with the proposed mechanism, because the acceleration needs to *consistently occur on the same segment of the sensory surface* (illustrated in Fig. 8). It is crucial for the process we propose that acceleration and skin location are thus correlated. As an analogy, consider a smudge on the glasses that moves with the head: the scenes change but the smudge remains, suggesting it is not a property of the world. Likewise, sudden accelerations attached to a particular skin segment suggest their cause is not in the world but in the receiving apparatus. Wiping the glasses removes the smudge, and a map change removes the acceleration. With disappearance of acceleration, any correlation between acceleration and location on the sensory surface also disappears. Thus achieved *decorrelation* is not an outcome of a teleological, goal-driven process, but a form of self-organization.

**Figure 8.**
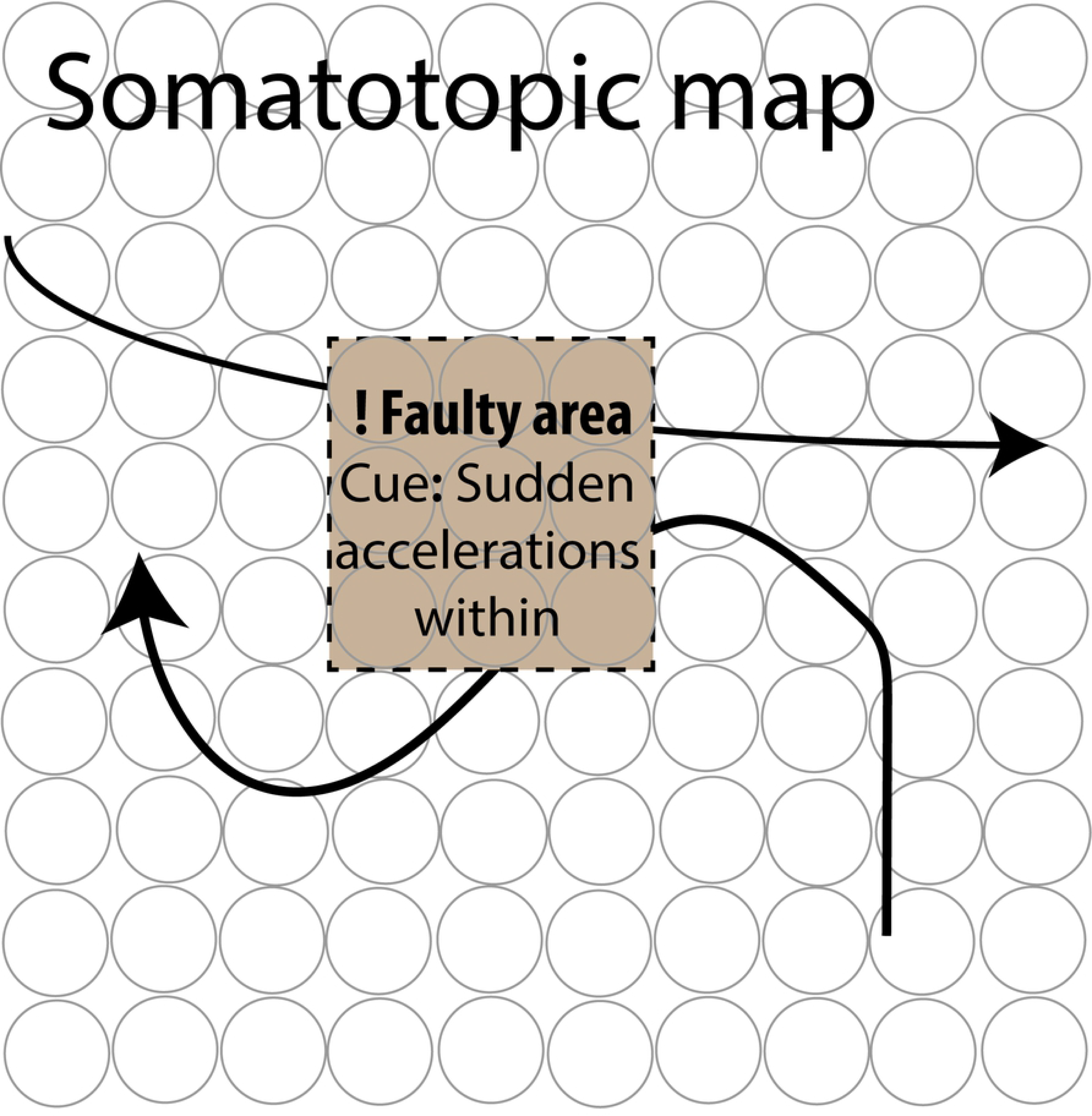
Schematic representation of our proposal that repeated, sudden accelerations that always start on the boundaries of a limited area of the sensory map signal an internal error in biological sensory maps, an example of which is the somatotopic map of the skin. The error may be a consequence of injury, surgical re-arrangement of the receptor surface, or any other cause that disrupts somatotopy. This learning principle may also be implemented in artificial neural networks. (The space-space schematic given here cannot represent velocity, but see Fig. 4A, Scrambled stimulus for an example of acceleration. Locations coloured red and green in that figure would fall within the shaded area here.)

The principle of decorrelation is one widely considered principle of efficient sensory coding. An insight from communication theory, applied to the brain, suggests that the correlated (redundant) messages impede transmission of sensory signals, wasting limited capacity of sensory channels (26). De- correlation is a proposed rectifying (or preventative) self-organizing process in neural populations [26-28]. It requires them to maintain similar levels of activity in all units (neurons), and minimize correlations, even when the stimulus distribution shifts or changes so that some values are overrepresented, or when faced with otherwise correlated stimuli. Example evidence in support of such a process are similar average responses and correlations observed in a population of orientation-sensitive neurons in cat V1 when exposed to different distributions of orientations, some of which were very biased [29].

On the other hand, the sensory system informs about the structure of the world, which is full of correlated features; thus it must preserve, explicitly store or make easily available information about the correlations [26].

The correlation of interest to us here is not between features of the distal stimulus (a physical object or event), but between *stimulus acceleration and its location* on the sensory surface – i.e., the features of the proximal stimulus (the way an object or event impacts the sensory surface). Map reorganization results in decorrelation and a better match with the world. Thus, the self-organizing principle thought to regulate neural population’s response to external stimulus features could also account for its self-correcting ability. A similar idea was put forward by New and Scholl [30] to account for motion-induced blindness (MIB). In MIB, a small object always falling on the *same segment* of the retina amidst a dynamic visual field quickly fades from awareness. They interpreted it as “the visual system’s attempt to separate distal stimuli from artifacts of damage to the visual system itself” (p. 655).

Neural mechanisms supporting the proposed changes likely involve context-sensitive, long-range connections between neurons in sensory maps and feedback from higher-order motion neurons on neurons that encode local motion and position. Involvement of the long-range connections allows the context of stimulation to disambiguate local input [31]. Filling-in of blind spots in vision and deafferented skin areas (numb spots) relies on such connections [31, 32] and blind spots are conceptually similar to our scrambled stimulus: both create discontinuities in the sensory input and both are ‘glued’ to a certain position on the receptor surface.

In summary, we propose a self-correcting system for maintenance of spatial maps, where the adjustment is triggered by accelerations recorded by the motion system whenever moving objects cross a particular location on the sensory surface, be it skin or retina. We describe this process as adaptation, defined operationally as a perceptual change that occurs following a relatively short exposure to sensory stimulation. We used an experimental paradigm for study of adaptation (the adapt-test cycle), and induce changes using seconds to minutes of exposure, as typically done in adaptation studies. Our dependent measure was perceptual response, and the proposed (beginning of) neural map change is hypothetical.

Other types of adaptation may also be relevant in an attempt to account for the effect we observed, and we turn to them now.

### C. Adaptation as reduced responsiveness (gain reduction) due to exposure to motion

Both scrambled and orderly patterns gave rise to perception of motion (see Fig. 7). Their repeated presentation would have activated motion-sensitive neurons, which have a strong presence in S1 [33-35], and presumably reduced their responsiveness. Adaptation effects are complex and occur at multiple levels (see [19] for review]. Neurons with receptive fields in the test area (affected by vibrators 3 and 4) would have adapted, as well as those in the near surround. Neurons at different levels in the somatosensory system would have adapted independently or as an inherited effect propagating from lower to higher levels. When adaptation is considered within this complex picture, we find at least two possible differences between our scrambled and orderly conditions.

First, *frequent direction change* might have resulted in *weaker* motion adaptation in the scrambled condition. Unlike the orderly pattern, in which direction of motion was constant during a single sweep – from elbow to wrist, or vice versa - in the scrambled pattern, direction change occurred twice during each sweep, in a double-u-turn. Perhaps direction-sensitive neurons at higher levels of processing stream, whose receptive fields cover the whole motion trajectory, adapted less when direction thus changed in the scrambled condition. This prediction seems contrary to our findings: we found worse performance – presumably indicating stronger adaptation – in the scrambled condition. Either the above factor did not contribute to different adaptation states in the two conditions, or other factors opposed it, creating the net effect we observed.

Second, *higher average speed* in the scrambled condition might have resulted in *stronger* adaptation in that condition. Unlike the orderly pattern, the scrambled pattern included accelerations during a single sweep: between locations 2 and 4, and 3 and 5, the speed was two times greater than speed used between other successive stimulations (the same ISOI of 120 ms was used for those pairs, which were 8 cm apart, as for the neighbouring locations that were 4 cm apart). We know from previous research that faster moving tactile stimuli create stronger adaptation [24, 25]. It is not clear how location-specific such an adaptation effect is, and whether our test stimulus moving between locations 3 and 4 would have tapped into it. If it had, it could be partly responsible for poorer direction discrimination in the scrambled condition.

We are currently unable to tease apart possible contributions of different adaptation effects described above.

### D. Simulated and actual rearrangements of receptor surface

We attempted to simulate location swap for skin patches under the middle two vibrators (3 and 4).

The simulation was constrained by the fact that vibration spread across the skin. We also used a relatively short period of conditioning (approximately 26 min per condition). Still, the manipulation induced a change in the way our healthy human observers perceived temporal order/motion between two skin locations.

Suppose that an actual surgery took place instead, swapping the locations of those skin patches, while preserving their innervation. Following recovery and a period of exposure to normal stimulation, we would expect the new spatial arrangement to be recognized by the brain, and normal spatial perception restored. Similar experiments, conducted on monkeys (study [2] described in the Introduction) represent classical demonstrations of plasticity of somatosensory maps (also see [6]).

Should our scrambled stimulus be applied for much longer time periods, and reinforced by other moving patterns that suggest a new skin arrangement (see Fig. 1), while other stimulation to the forearm is prevented, the outcome should be similar to an actual surgery. Motion direction (or temporal order) judgment should eventually be completely reversed, and there should also be errors in absolute localization, such that touch on a more distal skin patch should be mistakenly localized more proximally than touch on the patch proximal to it. These proposals are yet to be tested.

### E. Conclusion

In conclusion, a short exposure to a stimulation pattern that suggests a re-arrangement of the sensory surface results in an aftereffect consistent with the new arrangement, supporting the idea that motion plays a major role in organizing spatial maps in touch. Longer exposures and tests using a variety of sensory tasks would further test this idea and allow us to distinguish relative contributions of different adaptive processes. Our simulated-surgery paradigm is a potentially useful tool in the experimental study of plasticity in sensory maps in conscious humans.

## VI. ACKNOWLEDGMENTS

We thank Raymond Patton for constructing the vibrator array, and Timothy Turner and Diego Barneche for custom software.

